# Single-cell clonal lineage tracing identifies the transcriptional program controlling the cell fate decisions by neoantigen-specific CD8^+^ T cells

**DOI:** 10.1101/2025.08.13.670223

**Authors:** Ying Luo, Chen Yao, Tuoqi Wu

**Affiliations:** Department of Immunology, University of Texas Southwestern Medical Center, Dallas, TX; Kidney Cancer Program, Simmons Comprehensive Cancer Center, University of Texas Southwestern Medical Center, Dallas, TX; Simmons Comprehensive Cancer Center, University of Texas Southwestern Medical Center, Dallas, TX; Peter O’Donnell Jr. Brain Institute, University of Texas Southwestern Medical Center, Dallas, TX

## Abstract

Neoantigen-specific T cells specifically recognize tumor cells and are critical for cancer immunotherapies. However, the transcriptional program controlling the cell fate decisions by neoantigen-specific T cells is incompletely understood. Here, using joint single-cell transcriptome and TCR profiling, we mapped the clonal expansion and differentiation of neoantigen-specific CD8^+^ T cells in the tumor and draining lymph node in mouse prostate cancer. Compared to other antitumor CD8^+^ T cells and bystanders, neoantigen-specific CD8^+^ tumor-infiltrating lymphocytes (TILs) upregulated gene signatures of T cell activation and exhaustion. In the tumor draining lymph node, we identified TCF1^+^TOX^-^ T_SCM_, TCF1^+^TOX^+^ T_PEX_, and TCF1^-^TOX^+^ effector-like T_EX_ subsets among neoantigen-specific CD8^+^ T cells. Clonal tracing analysis of neoantigen-specific CD8^+^ T cells revealed greater clonal expansion in divergent clones and less expansion in clones biased towards T_EX,_ T_PEX_, or T_SCM_. The T_PEX_ subset had greater clonal diversity and likely represented the root of neoantigen-specific CD8^+^ T cell differentiation, whereas highly clonally expanded effector-like T_EX_ cells were positioned at the branch point where neoantigen-specific clones exited the lymph node and differentiated into T_EX_ TILs. Notably, T_SCM_ differentiation of neoantigen-specific CD8^+^ clones in the lymph node negatively correlated with exhaustion and clonal expansion of the same clones in the tumor. In addition, the gene signature of neoantigen-specific clones biased toward tumor infiltration relative to lymph node residence predicted a poorer response to immune checkpoint inhibitor. Together, we identified the transcriptional program that controls the cell fate choices by neoantigen-specific CD8^+^ T cells and correlates with clinical outcomes in cancer patients.

## Introduction

CD8^+^ T cells are critical for the immune protection against malignancies and pathogens. Antigen recognition activates naïve CD8^+^ T cells and triggers effector differentiation and clonal expansion. Effector CD8^+^ T cells are heterogeneous and contain short-lived effector cells and memory precursor effector cells^1–3^. After antigen clearance, surviving memory precursor effector cells develop into memory T cells which persist independent of antigen and mediate long-term protection against the same antigen ^4,5^. However, antigen is not cleared in chronic viral infection or cancer. Persist antigen stimulation diverges CD8^+^ T cells from their normal naïve→effector→memory path into an alternative cell fate, termed exhaustion ^6,7^. Exhausted CD8^+^ T (T_EX_) cells upregulate immune checkpoints such as PD1, CTLA4, TIM3, LAG3, and TIGIT and have reduced proliferation and effector function ^6,7^. Immune checkpoint inhibitors reinvigorate the immunity of T_EX_ cells and have shown remarkable efficacy in a subset of cancer patients ^8–10^.

It is now known that T_EX_ cells are heterogeneous. The expression of transcription factor TCF1 separates T_EX_ cells into two populations. The TCF1^-^ terminally exhausted subset is short-lived and expressed higher levels of immune checkpoints, whereas the TCF1^+^ stem cell-like subset maintains the T_EX_ pool through self-renewal and repopulating the terminally exhausted subset ^11–21^. TCF1^+^ stem-like T cells, also termed progenitor exhausted T cells (T_PEX_), promote the efficacy of immune checkpoint inhibitors (ICIs) and chimeric antigen receptor (CAR) T cell therapy ^15–18,22–25^. The differentiation of T_PEX_ cells is driven by transcription factors such as TCF1, BCL6, BACH2, TOX, FOXP1, FOXO1, and MYB and repressed by BLIMP1, IRF4, NRF2, and type I interferon ^11–15,19,26–37^. T_PEX_ cells are strategically positioned in lymphoid tissues in both cancer and chronic viral infection and function as a reservoir to maintain T-cell immunity and respond to immune checkpoint inhibitors ^12–14,21,38–42^. In addition to terminally exhausted T cells (term T_EX_), T_PEX_ cells also differentiate into other lineages including an effector-like subset (eff- like T_EX_) that displays greater cytotoxicity and promotes control of viruses and tumor cells ^43–47^. Eff-like T_EX_ differentiation relies on transcription factors KLF2 and BATF and IL21 derived from CD4^+^ T cells^37,43,48^. Single-cell RNA-sequencing (scRNA-seq) identified various exhaustion states of tumor infiltrating T cells (TILs) including those recognizing neoantigens ^49–53^. However, the heterogeneous differentiation states and lineage relationship of neoantigen-specific CD8^+^ T cells in the tumor and lymphoid tissue is incompletely understood.

In this study, we performed joint single-cell transcriptome and TCR sequencing (scRNA+TCR-seq) with CD8^+^ T cells responding to a syngeneic prostate cancer model with a well-defined neoepitope in SPAS-1 naturally generated during tumorigenesis ^54^. Using IFNγ-YFP reporter (GREAT) mice^55^, we found that neoantigen SPAS-1-specific CD8^+^ TILs as well as a subset of SPAS-1 tetramer^-^ antitumor TILs showed a predominant T_EX_ phenotype, expressed both IFNγ and PD1 and clonally expanded whereas bystander TILs did not express IFNγ or PD1 and showed limited clonal expansion. In the tumor-draining lymph node, neoantigen-specific CD8^+^ T cells consisted of both PD1^+^ (eff-like T_EX_, T_PEX_) and PD1^-^ (T_SCM_) subsets. The response against neoantigen SPAS-1 was dominated by a few TCR clones present in both the lymph node and tumor, allowing us to perform clonal tracing. The divergent clones accounted for the majority of neoantigen-specific CD8^+^ T cells and were more clonally expanded, whereas lineage-biased clones were less clonally expanded. The T_PEX_ subset showed the highest TCR diversity among neoantigen-specific CD8^+^ subsets in the lymph node and represented the root subset in the differentiation trajectory of neoantigen-specific CD8^+^ T cells. The eff-like T_EX_ subset was the most clonally expanded subset and may represent a branch point where neoantigen-specific CD8^+^ T cells exited the lymph node and infiltrated the tumor. We demonstrated that T_SCM_ bias in the neoantigen-specific CD8^+^ clones in the lymph node correlated with reduced exhaustion and less clonal expansion in the tumor. Lastly, we showed that the gene signature of tumor-biased neoantigen-specific CD8^+^ clones positively correlated with overall survival but negatively correlated with the ICI-induced response in cancer patients. Together, our results uncover the transcriptional program among neoantigen-specific CD8^+^ clones responding to prostate tumor that influences their cell fate decisions and correlates with clinical outcome in patients.

## Results

### Antigen recognition and tissue environment dictate the transcriptional program of CD8^+^ T cells responding to prostate cancer

To investigate the CD8^+^ T cell response against neoantigen, we used a syngeneic mouse prostate tumor TRAMP-C2 model, which contains an immunodominant neoantigen, SPAS-1, derived from a point mutation ^54^. We inoculated TRAMP-C2 to IFNγ-YFP reporter (GREAT) mice^55^ to monitor antitumor T cells through the expression of IFNγ. On day 30 post-tumor inoculation, neoantigen-specific CD8^+^ T cells were identified by the SPAS-1 tetramer in the tumor and draining lymph node. Tumor-infiltrating SPAS-1 specific CD8^+^ T cells were IFNγ-YFP^+^ PD1^+^ (**Figure 1A**), suggesting that neoantigen-specific CD8^+^ T cells were producing IFNγ in response to antigen recognition in the tumor. Within the SPAS-1 tetramer^-^ CD8^+^ tumor-infiltrating lymphocytes (TILs), there were IFNγ-YFP^+^ PD1^+^ and IFNγ-YFP^-^ PD1^-^ subsets (**Figure 1A**). To comprehensively understand the transcriptional programs of neoantigen-specific CD8^+^ T cells and other CD8^+^ TILs, we performed a scRNA+TCR-seq experiment with SPAS-1 tetramer^+^ CD8^+^ T cells in the tumor and draining lymph node as well as IFNγ-YFP^+^ PD1^+^ and IFNγ-YFP^-^ PD1^-^ CD8^+^ TILs (**Figure 1B**). We obtained scRNA-seq profiles from 9,656 cells across four samples. An average of 6,769 mRNA molecules and 2,242 genes were detected in each cell.

**Figure 1.**
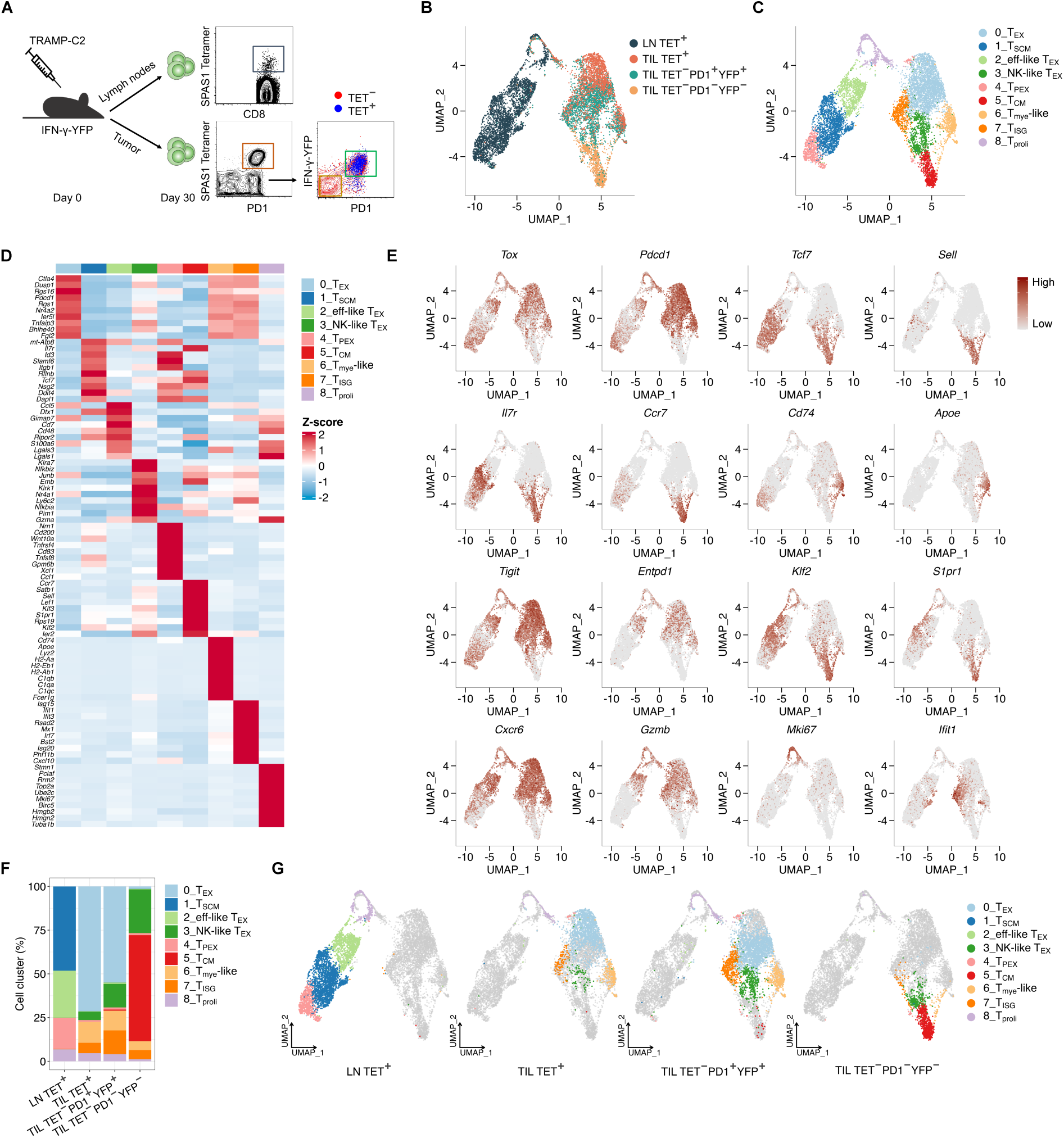
Antigen recognition and tissue environment control the transcriptional program of CD8^+^ T cells responding to prostate cancer. (**A**) Schematic illustration of the experimental design and cell sorting strategy. SPAS-1 tetramer (Tet)^+^ CD8^+^ T cells were collected from tumor-draining lymph nodes (LN) and tumor-infiltrating lymphocyte (TIL). SPAS-1 tetramer^-^ TILs were sorted into PD1^+^IFNγ-YFP^+^ and PD1^-^IFNγ-YFP^-^ subsets. (**B**) UMAP visualization of the single-cell RNA-sequencing (scRNA-seq) portion of the scRNA+TCR-seq data combined from LN Tet^+^, TIL Tet^+^, TIL Tet^-^PD1^+^IFNγ-YFP^+^, and TIL Tet^-^PD1^-^IFNγ-YFP^-^ populations at day 30 post-TRAMP-C2 inoculation. Each dot represents a single cell, colored by sample identity. (**C**) UMAP plot of the scRNA-seq portion of the scRNA+TCR-seq data combined from LN Tet^+^, TIL Tet^+^, TIL Tet^-^PD1^+^IFNγ-YFP^+^, and TIL Tet^-^PD1^-^IFNγ-YFP^-^ cells at day 30 post-TRAMP-C2 inoculation. Nine annotated T cell subsets are color-coded, with each dot representing a single cell. (**D**) Heatmap showing the expression of key marker genes across 9 annotated T cell subsets as defined in panel C, with each column corresponding to a T cell subset. The color scale represents z-score-transformed transcript levels for each gene. (**E**) UMAP visualization of single-cell transcript levels for the selected genes. Expression levels are color-coded with dark red indicating high expression. (**F**) Stacked bar plot depicting the relative abundance of each annotated T cell subsets across different samples. (**G**) UMAP distribution of the annotated T cell subsets within each sample.

Uniform manifold approximation and projection (UMAP) was performed with scRNA-seq profiles for dimension reduction (**Figure 1B**). Lymph node (LN) SPAS-1 tetramer^+^ CD8^+^ T cells occupied an area in the UMAP distant from SPAS-1 tetramer^+^ TILs, suggesting a profound impact of tumor microenvironment on the transcriptional program (**Figure 1B**). While tetramer^-^IFNγ-YFP^+^ PD1^+^ TILs showed a partial overlap with SPAS-1 tetramer^+^ TILs in the UMAP, IFNγ-YFP^-^ PD1^-^ TILs did not comingle with tetramer^+^ TILs (**Figure 1B**). This result suggests that YFP^+^ PD1^+^ TILs are transcriptionally more similar to neoantigen-specific TILs than YFP^-^ PD1^-^ TILs likely as a result of recognizing other tumor antigens. We next performed an unsupervised clustering which separated CD8^+^ T cells into nine clusters (**Figure 1C**). The largest cluster in TILs was exhausted CD8^+^ T cells (T_EX_) that upregulated chemokine receptor *Cxcr6*, pro-exhaustion transcription factors *Tox* and *Nr4a2* as well as immune checkpoints including *Ctla4*, *Pdcd1* (PD1), *Tigit*, and *Entpd1* (CD39) (**Figure 1C-E**; **Figure S1A**). The T_EX_ subset consisted of SPAS-1 tetramer^+^ TILs and tetramer^-^IFNγ-YFP^+^PD1^+^ TILs (**Figure 1C,F**). A central memory (T_CM_) TIL subset that expressed high levels of *Sell* (CD62L), *Ccr7*, *Tcf7* and *Klf2* was composed of IFNγ-YFP^-^ PD1^-^ TILs (**Figure 1C-G**). We identified a natural killer (NK)-like TIL subset expressing multiple NK receptors including *Klrk1* (NKG2D) (**Figure 1C,D**). In addition, we observed a TIL subset expressing interferon-stimulated genes (ISG) and a TIL subset expressing myeloid markers including *Cd74*, *Apoe* and *Lyz2*, which mostly consisted of SPAS-1 tetramer^+^ TILs and tetramer^-^IFNγ-YFP^+^ PD1^+^ TILs (**Figure 1C-G**). A subset of proliferating cells was found in SPAS-1 tetramer^+^ TILs, IFNγ-YFP^+^ PD1^+^ TILs and LN SPAS-1 tetramer^+^ CD8^+^ T cells but not in IFNγ-YFP^-^ PD1^-^ TILs (**Figure 1B,C,G**).

There were four clusters among LN SPAS-1 tetramer^+^ CD8^+^ T cells (**Figure 1C**). Besides proliferating cells, the other three clusters showed distinct expression pattern of genes associated with stem/memory T cells or exhausted T cells. One LN SPAS-1 tetramer^+^ subset expressed high levels of stem-like T cell markers including *Tcf7*, *Il7r*, *Slamf6*, and *Id3*, intermediate levels of T_CM_ markers including *Ccf7* and *Sell*, and low levels of genes associated with cytotoxicity or exhaustion such as *Tox* (**Figure 1C-F**; **Figure S1A**). This *Tcf7*^+^*Tox*^-^*Pdcd1*^-^T stem cell memory (T_SCM_) subset was previously shown to be the bona fide responder to PD1/PD-L1 blockade ^56^. Conversely, an effector-like exhausted subset (eff-like T_EX_) in LN SPAS-1 tetramer^+^ CD8^+^ T cells downregulated stem/memory T cell markers including *Tcf7* and upregulated genes encoding cytotoxic proteins (*Gzmb*) and exhaustion-related genes (*Tox*, *Pdcd1*, *Tigit, Entpd1*) (**Figure 1C-F**). LN SPAS-1 tetramer^+^ CD8^+^ T cells also contained a progenitor exhausted subset (T_PEX_) that co-expressed stem-like T cell markers (*Tcf7*, *Slamf6*, *Id3*) and exhaustion-related genes (*Tox*, *Pdcd1*, *Tigit*) (**Figure 1C-F**; **Figure S1A**). Unlike eff-like T_EX_ cells, T_PEX_ cells did not express *Entpd1* or cytotoxic molecules. T_PEX_ cells expressed lower level of *Klf2*, *S1pr1* and *Sell* than T_SCM_ cells, suggesting a potential difference in the ability to enter circulation. Together, our data show that both antigen recognition and tissue environment determine the transcriptional program of CD8^+^ T cells responding to prostate cancer.

### scRNA-seq identifies the unique transcriptional program of neoantigen-specific CD8^+^ TILs

We have demonstrated that both SPAS-1 tetramer^+^ CD8^+^ TILs and tetramer^-^IFNγ-YFP^+^ PD1^+^ CD8^+^ TILs produced IFNγ and proliferated in response to prostate cancer. However, these two populations did not fully comingle in the UMAP especially within the T_EX_ subset (**Figure 2A,B**), suggesting different transcriptional programming. Compared to tetramer^-^IFNγ-YFP^+^ PD1^+^ TILs, neoantigen-specific TILs upregulated both costimulatory receptors such as *Tnfrsf9* (4-1BB) and *Tnfrsf4* (OX40) and immune checkpoints such as *Lag3* and *Tigit* (**Figure 2C**). Gene-set enrichment analysis (GSEA) showed that neoantigen-specific TILs upregulated gene signatures associated with T-cell activation, proliferation, and cytokine production and downregulated gene signatures associated with interferon response and ribosome (**Figure 2D**). Therefore, compared to other antitumor CD8^+^ TILs, neoantigen-specific CD8^+^ TILs showed a hybrid phenotype of T cell activation and exhaustion.

**Figure 2.**
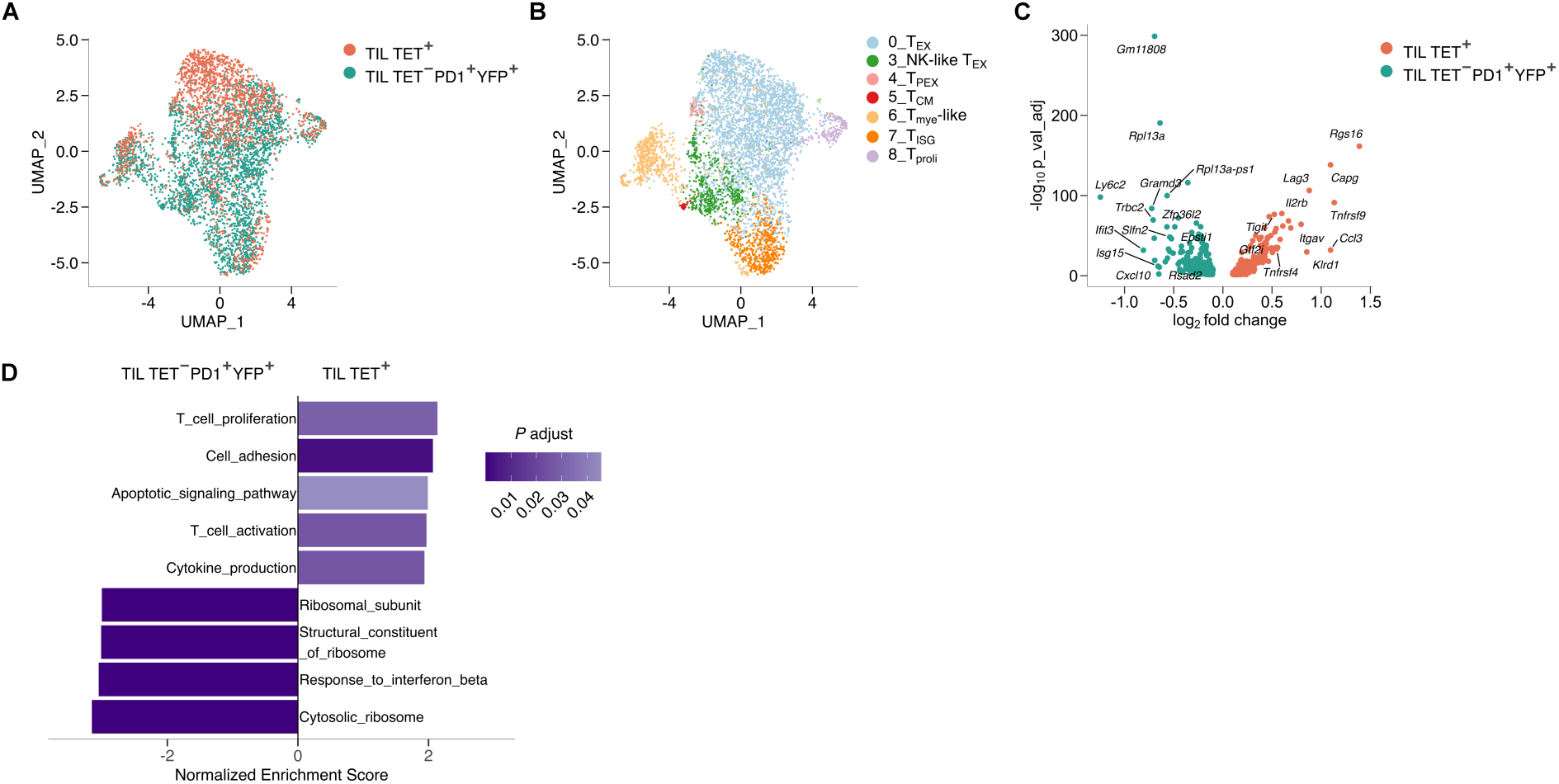
Neoantigen-specific CD8^+^ T cells show a hybrid phenotype of T cell activation and exhaustion. (**A**) UMAP plot of the scRNA-seq component of the integrated scRNA+TCR-seq data from SPAS-1 tetramer (Tet)^+^ CD8^+^ TILs and Tet^-^PD1^+^IFNγ-YFP^+^ CD8^+^ TILs at day 30 post- TRAMP-C2 inoculation. Each dot represents a single cell, colored by sample identity. (**B**) UMAP plot displaying the scRNA-seq portion of the scRNA+TCR-seq data combined from Tet^+^ TILs and Tet^-^PD1^+^IFNγ-YFP^+^ TILs at day 30 post-TRAMP-C2 inoculation, with cells colored by the annotated T cell subsets identified with Seurat. (**C**) Volcano plot showing differentially expressed genes between Tet^+^ TILs and Tet^-^PD1^+^IFNγ-YFP^+^ TILs. (**D**) Bar graph showing the differentially enriched pathways identified by gene set enrichment analysis (GSEA) comparing Tet^+^ TILs and Tet^-^PD1^+^IFNγ-YFP^+^ TILs at day 30 post-TRAMP-C2 inoculation. The x-axis represents the normalized enrichment score for each pathway, and the y-axis shows the corresponding pathways. The color-code indicates the adjusted *P* value for each pathway.

### The response against neoantigen is dominated by a few TCR clones

Next, we sought to perform TCR clonal analysis of neoantigen SPAS-1 specific CD8^+^ T cells in the lymph node and TIL as well as IFNγ-YFP^+^ PD1^+^ and IFNγ-YFP^-^ PD1^-^ TILs. Paired TCRα and TCRβ sequences were detected from 83% of the scRNA+TCR-seq profiles described above. We identified 1,286 T cell clones, defined by T cells with identical amino acid sequences at the CDR3 regions of TCRα and TCRβ. A total of 192 expanded T cell clones (>1 cell per clone) were identified. Among SPAS-1 tetramer^+^ CD8^+^ T cells in the lymph node and tumor, 98.87% and 99.59% cells were expanded T cell clones, respectively, indicating extensive clonal expansion (**Figure 3A,B**). 91.98% tetramer^-^ IFNγ-YFP^+^ PD1^+^ CD8^+^ T cells were expanded clones, whereas only 8.46% tetramer^-^ IFNγ-YFP^-^ PD1^-^ CD8^+^ T cells clonally expanded (**Figure 3A,B**). Thus, the majority of tetramer^-^ IFNγ-YFP^-^ PD1^-^ cells are likely bystander T cells attracted to the tumor. We categorized the size of T cell clones into large (>= 523 cells per clone), medium (46-522 cells per clone), small (2-45 cells per clone), and single (1 cell per clone) (**Figure 3A,B**). Notably, more than 50% of SPAS-1 specific TILs were large clones, whereas large clones accounted for nearly 50% of SPAS-1 specific LN CD8^+^ T cells and IFNγ-YFP^+^ PD1^+^ TILs (**Figure 3A-C**). The top 10 clones constituted ∼88.22% of SPAS-1 specific TILs (**Figure S2A**), which had a lower Shannon diversity compared to LN SPAS-1 specific CD8^+^ T cells, IFNγ-YFP^+^ PD1^+^ TILs, and IFNγ-YFP^-^ PD1^-^ TILs (**Figure 3D**). Chord diagrams of TCR V and J pairing showed clonal dominance by a single TCRβ V-J pair in neoantigen SPAS-1 specific CD8^+^ T cells in both the tumor and lymph node (**Figure 3E**). Thus, our TCR clonality analysis demonstrates that the CD8^+^ T cell response against neoantigen is heavily skewed towards a few dominant TCR clones and that combination of PD1 and IFNγ expression distinguishes tumor antigen-specific CD8^+^ T cells from bystanders.

**Figure 3.**
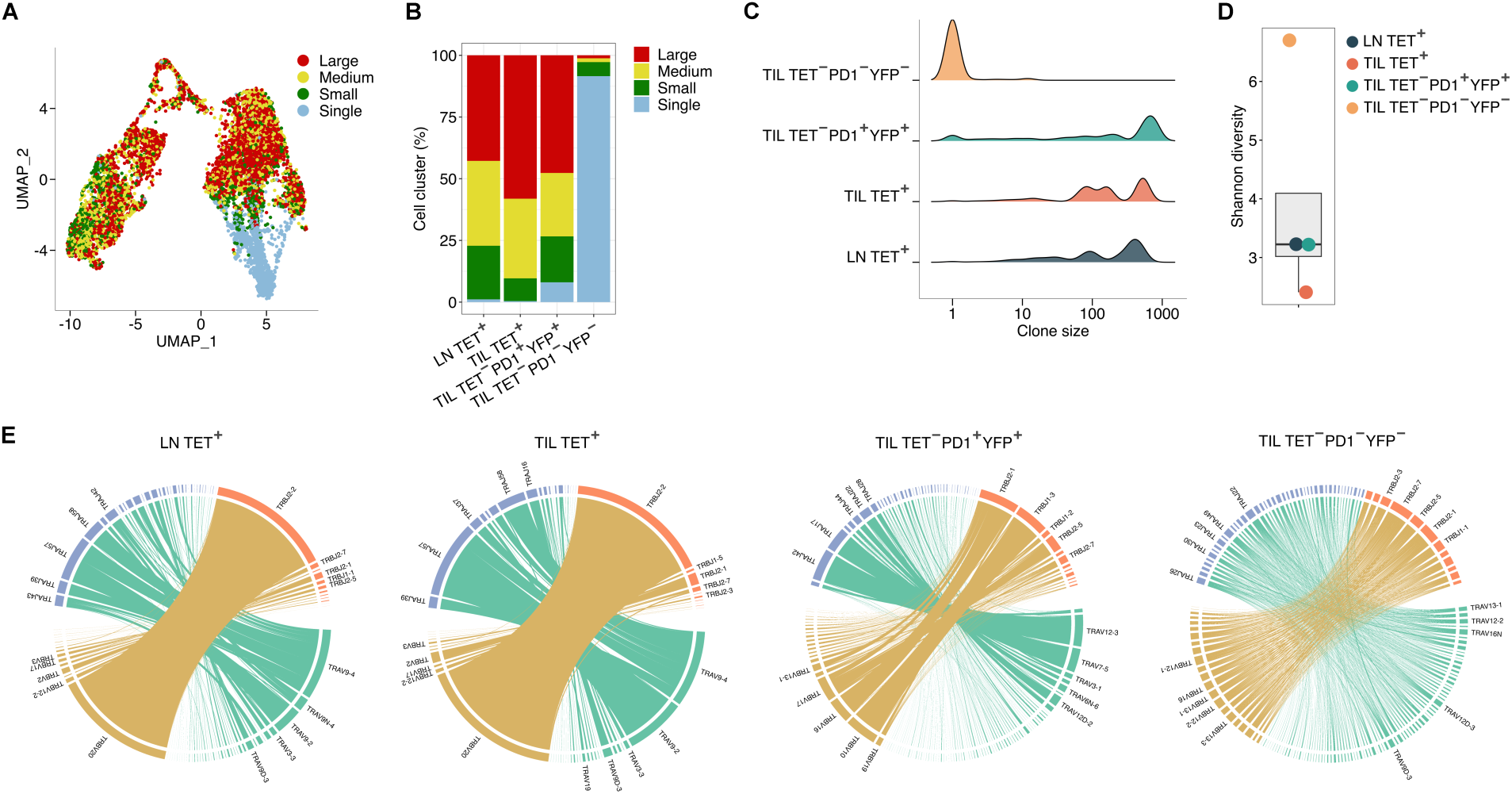
Neoantigen-specific CD8^+^ T cells show strong clonal dominance. (**A**) UMAP plot showing single cell embedding color-coded by clonal size: large (>=523 cells), medium (46-522 cells), small (2-45 cells), and single (1 cell). Each dot represents an individual cell, with color indicating the clonal size category. (**B**) Stacked bar plot quantifying the relative proportions of clones classified by the size including large (>=523 cells), medium (46-522 cells), small (2-45 cells), and single (1 cell) across samples. (**C**) Ridge plots showing the distribution of clone sizes across samples. (**D**) Box plot illustrating clonal diversity across populations, as measured by the Shannon diversity index. (**E**) Chord diagrams showing the pairing of TCR α and β chain variable (V) and joining (J) genes. Each band represents a unique V-J pair, with band thickness corresponding to the frequency of each unique V-J pair in the respective samples.

### Clone tracing uncovers distinct differentiation fates of neoantigen-specific CD8^+^ T cells

To determine the relationship between TCR clonality and differentiation, we performed a clonotypic analysis in each CD8^+^ subset (**Figure 4A**). Eff-like T_EX_ CD8^+^ T cells in the lymph node had the highest percentage of large T cell clones followed by T_EX_ CD8^+^ TILs and proliferating cells (**Figure 4A**, **S3A**). T_CM_ TIL, which were primarily IFNγ-YFP^-^ PD1^-^ bystanders, consisted of single T cell clones (**Figure 4A**, **S3A**). Notably, although neoantigen-specific T_PEX_ cells were predominantly expanded clones, the clone size of T_PEX_ was smaller than their T_SCM_ and eff-like T_EX_ counterparts in the lymph node (**Figure 4A,B**). Consistently, Shannon diversity was the lowest in eff-like T_EX_ CD8^+^ T cells followed by T_EX_, proliferating, and T_SCM_ subsets, whereas the T_PEX_ subset had the highest clonal diversity among lymph node CD8^+^ subsets (**Figure 4C**). Thus, clonal heterogeneity among tumor-specific CD8^+^ T cells is lowest in eff-like T_EX_ subset and high in the T_PEX_ subset.

**Figure 4.**
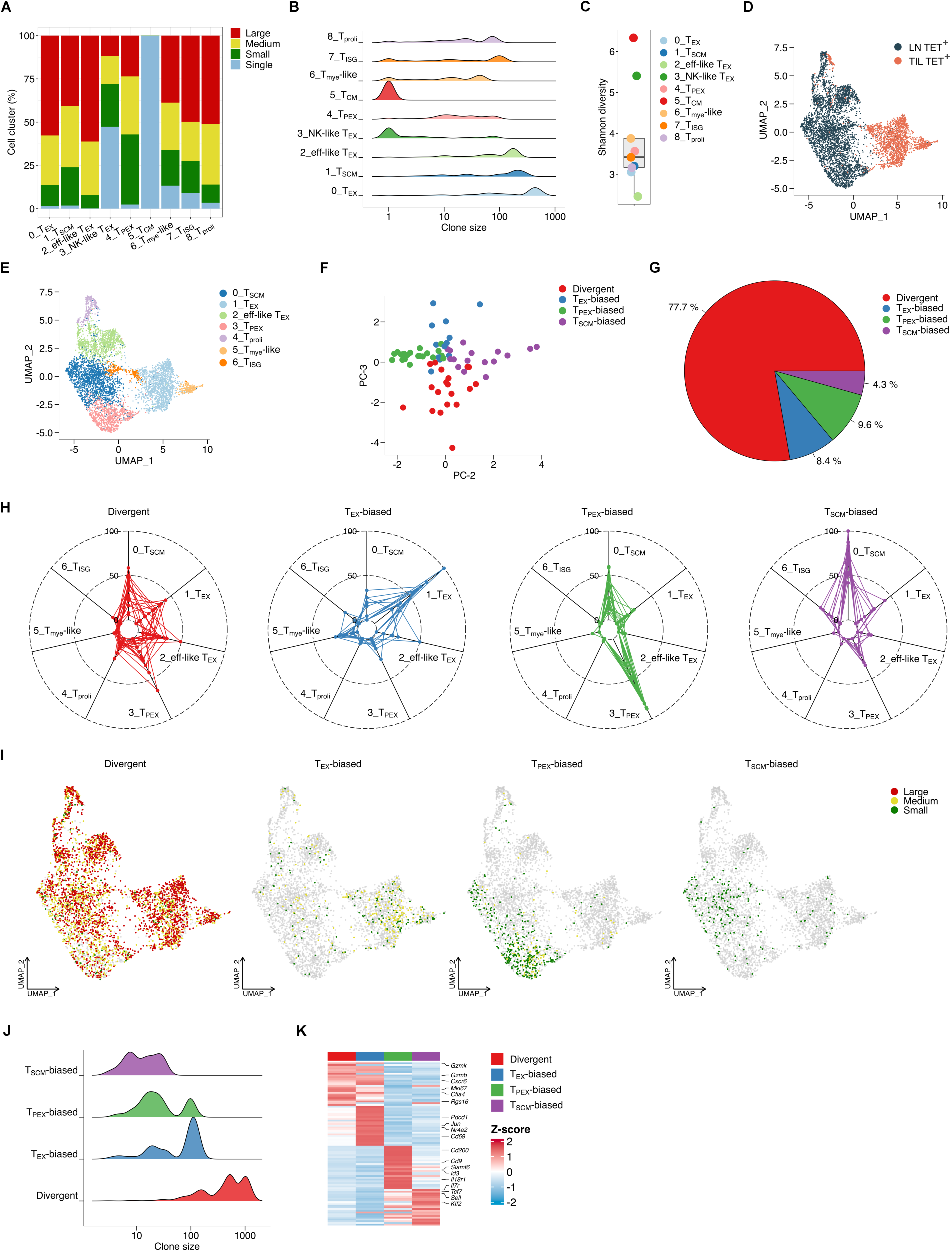
Clonal tracing reveals lineage preference in neoantigen-specific CD8^+^ T cells. (**A**) Stacked bar plot illustrating the relative proportions of large (>=523 cells), medium (46-522 cells), small (2-45 cells), and single (1 cell) clones across the annotated T cell subsets. (**B**) Ridge plots showing the distribution of clone sizes across the annotated T cell subsets. (**C**) Box plot illustrating the clonal diversity across the annotated T cell subsets, as quantified by the Shannon diversity index. (**D**) UMAP plot of the scRNA-seq portion of the scRNA+TCR-seq data combined from lymph node (LN) SPAS-1 tetramer (Tet)^+^ CD8^+^ T cells and Tet^+^ CD8^+^ TILs at day 30 post-TRAMP-C2 inoculation. Cells are color-coded by population identity. (**E**) UMAP plot of the scRNA-seq portion of the scRNA+TCR-seq data combined from LN Tet^+^ CD8^+^ T cells and Tet^+^ CD8^+^ TILs at day 30 post-TRAMP-C2 inoculation, with each cell colored according to the 7 annotated T cell subsets identified using Seurat. (**F**) A principal component analysis (PCA) plot of divergent, T_EX_-biased, T_PEX_-biased, and T_SCM_-biased clones. (**G**) Pie chart showing the relative cell number proportions of divergent, T_EX_-biased, T_PEX_-biased, and T_SCM_-biased clones within Tet^+^ CD8^+^ T cells. (**H**) Radar plots depicting the percentages of the annotated T cell subsets within divergent, T_EX_-biased, T_PEX_-biased, and T_SCM_-biased clones. (**I**) UMAP plots showing cell embeddings of divergent, T_EX_-biased, T_PEX_-biased, and T_SCM_-biased clones, colored by large, medium and small clone sizes defined in Panel A. (**J**) Ridge plots showing the distribution of clone sizes among divergent, T_EX_-biased, T_PEX_-biased, and T_SCM_-biased clones. (**K**) Heatmap showing clone-pattern-specific genes in divergent, T_EX_-biased, T_PEX_-biased, and T_SCM_-biased clones.

Next, we sought to evaluate potential differentiation bias among different TCR clones of neoantigen-specific CD8^+^ T cells. Dimension reduction and clustering were performed with SPAS-1 tetramer^+^ CD8^+^ T cells in the lymph node and tumor (**Figure 4D,E**). By clustering TCR clones based on their distribution in different T cell subsets, we identified four clonal differentiation patterns (**Figure 4F**). Clones corresponding to 77.7% of SPAS-1 specific CD8^+^ T cells showed a divergent phenotype and were similarly distributed among major differentiation fates including T_EX_, T_SCM_, eff-like T_EX_, T_PEX_ and proliferating subsets (**Figure 4G,H**). Because a TCR clonotype is theoretically derived from a single naïve precursor cell, our data suggest that a large proportion of naïve neoantigen-specific CD8^+^ T cells respond to prostate tumor in a “one cell, multiple fate” manner. The second largest (9.6%) clonal behavior pattern contained neoantigen-specific CD8^+^ T cells that showed a strong bias towards differentiation of T_PEX_ cells (**Figure 4G,H**). We also identified TCR clones in SPAS-1 specific CD8^+^ T cells that skewed towards differentiation of T_EX_ cells (8.4%) or T_SCM_ cells (4.3%) (**Figure 4G,H**). The divergent clones underwent the highest clonal expansion and had an average of 168 cells per clone (**Figure 4I,J**). Notably, T_SCM_-biased clones (10 cells per clone), T_EX_-biased clones (28 cells per clone), and T_PEX_-biased clones (18 cells per clone) showed lower clonal expansion and consisted of medium and small clones (**Figure 4I,J**). Notably, divergent clones upregulated genes associated with effector function (*Gzmk*, *Gzmb*) and exhaustion (*Ctla4*) as well as pathways such as cell cycle, cell death, and cytokine production (**Figure 4K**, **S3B**). T_SCM_-biased clones increased expression of memory marker genes including *Il7r* and *Sell* and upregulated ribosomal protein, T cell activation, mRNA binding, and chemokine binding pathways (**Figure 4K**, **S3B**). Together, these results reveal that neoantigen-specific CD8^+^ T cells contain highly clonally expanded divergent and less expanded T_SCM_-biased, T_EX_-biased and T_PEX_-biased clones.

### Pseudotime analysis reveals the differentiation trajectory of neoantigen-specific CD8^+^ T cells

To determine the differentiation trajectories of neoantigen-specific CD8^+^ T cells, we performed a differentiation trajectory and pseudotime analysis of SPAS-1 specific CD8^+^ T cells in the tumor and lymph node (**Figure 5A-D**). The trajectory analysis revealed an overall differentiation from SPAS-1 specific CD8^+^ T cells in the lymph node towards those in the tumor (**Figure 5A**). SPAS-1 specific CD8^+^ T cells contained seven major clusters including T_SCM_, T_EX_, eff-like T_EX_, T_PEX_, proliferating, ISG, and myeloid-like subsets (**Figure 4E**). The root of the differentiation trajectory was formed by predominantly T_PEX_ cells (**Figure 5B,C**). The first branch corresponded to a T_PEX_→ T_SCM_ → eff-like T_EX_ differentiation trajectory (**Figure 5B-D**). At the branch point, eff-like T_EX_ CD8^+^ T cells diverged into the proliferating subset or tumor-infiltrating subsets including the T_EX_ subset, the ISG subset and the myeloid-like subset (**Figure 5B-D**). We observed several distinct gene expression patterns along the differentiation trajectory of neoantigen-specific CD8^+^ T cells (**Figure 5E**). *Tcf7* was downregulated during T_SCM_ → eff-like T_EX_ transition and was maintained in a low level among TIL subsets (**Figure 5E**). *Tox* and immune checkpoints including *Pdcd1*, *Tigit*, *Ctla4*, and *Lag3* were expressed in T_PEX_ cells, downregulated during T_PEX_ → T_SCM_ transition, upregulated in eff-like T_EX_ cells and either maintained in a similar level or further upregulated after differentiation into TIL T_EX_ subsets (**Figure 5E**). Other genes such as *Havcr2* (TIM3), *Gzmb*, and *Cxcr6* were upregulated during T_SCM_ → eff-like T_EX_ transition and maintained in high levels in TIL T_EX_ subsets (**Figure 5E**). Thus, pseudotime analysis demonstrated the eff- like T_EX_ subset as a key branch point during the differentiation from the T_PEX_ subset to tumor-infiltrating T_EX_ subsets.

**Figure 5.**
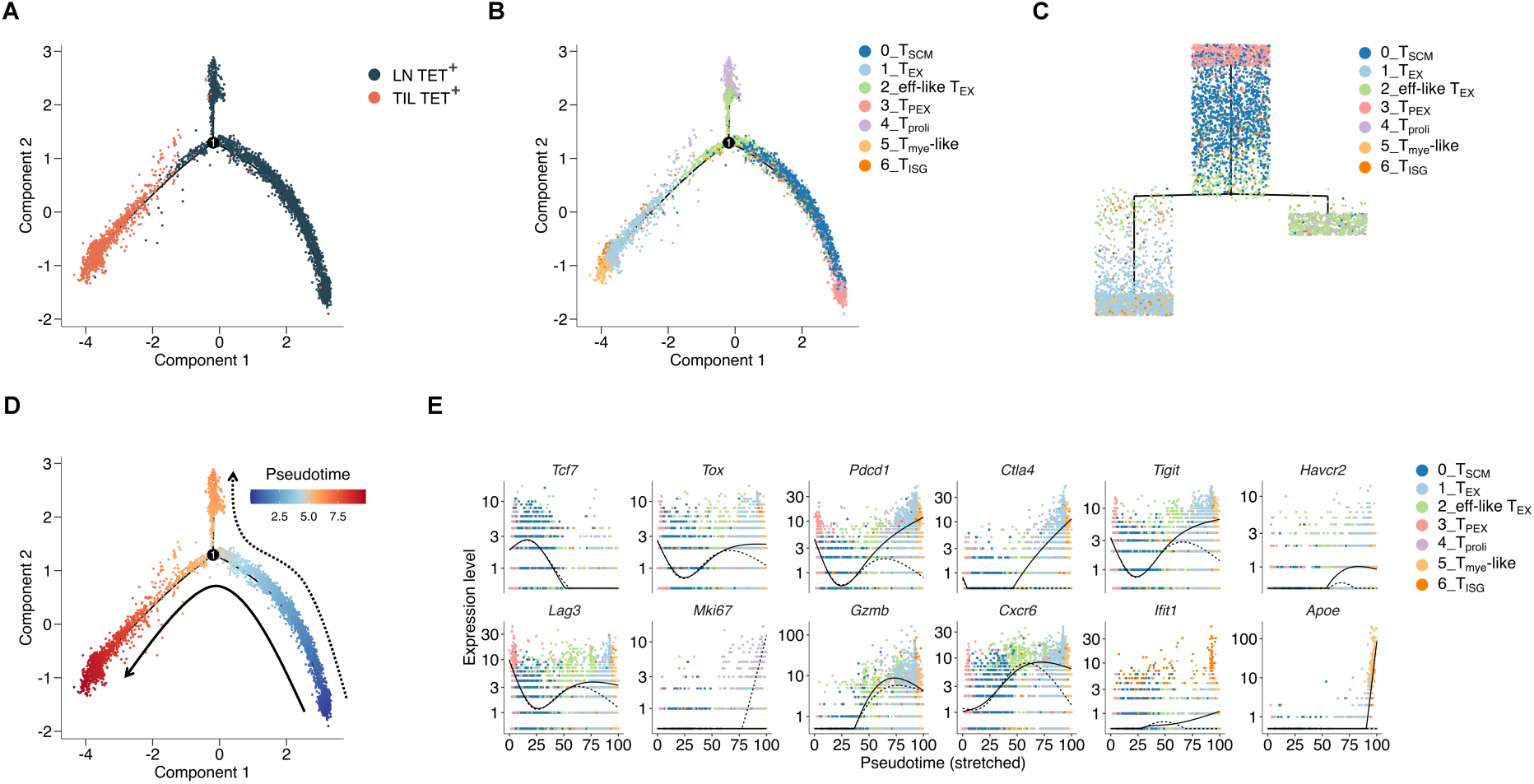
Pseudotime analysis of neoantigen-specific CD8^+^ T cells reveals distinct differentiation trajectories. (**A**) Single-cell trajectory plot generated by Monocle 2 showing lineage relationships among lymph node (LN) SPAS-1 tetramer (Tet)^+^ CD8^+^ T cells and Tet^+^ CD8^+^ TILs. Cells are colored by sample identity. (**B**) Single-cell trajectory plot generated by Monocle 2 showing lineage relationships among the annotated T cell subsets. Cells are colored by T cell subset identity. (**C**) Monocle trajectory tree plot generated by Monocle 2 illustrating transcriptional progression among the annotated T cell subsets. Cells are colored by T cell subset identity. (**D**) Single-cell trajectory plot generated by Monocle 2 showing predicted cell differentiation along pseudotime. Cells are colored by pseudotime values. (**E**) Plots showing gene expression dynamics over pseudotime. Solid and dash lines denote T_PEX_ → T_SCM_ → eff-like T_EX_→ T_EX_ trajectory and T_PEX_ → T_SCM_ → eff-like T_EX_→ T_proli_ trajectory, respectively. Colors denote the annotated T cell subset identity.

### A lineage preference towards T_SCM_ in the lymph node imprints a distinct cell fate in the tumor

We aimed to understand how differentiation of neoantigen-specific CD8^+^ T cells in the lymph node impacts their cell fate in the tumor through clonal lineage tracing. We first categorized neoantigen-specific CD8^+^ T cell clones based on their clonal expansion in the tumor. Notably, high clonal expansion in the tumor correlated with a lower frequency of T_SCM_ cells and a higher frequency of eff-like T_EX_ cells of the same clones in the lymph node (**Figure 6A**). Consistently, neoantigen-specific T cell clones with high expansion in the tumor showed a higher level of effector molecule *Gzmk* and a lower level of memory marker *Sell* than clones with low expansion in the tumor (**Figure 6B**). Next, we used the exhaustion score of TILs to categorize neoantigen-specific CD8^+^ T cell clones (**Figure 6C**). Neoantigen-specific T cell clones with a lower exhaustion score in the tumor had a higher frequency of the T_SCM_ subset and lower frequencies of the eff-like T_EX_ and T_PEX_ subsets in the lymph node than clones with a higher exhaustion score (**Figure 6D**). Compared to high exhaustion clones, low exhaustion clones upregulated markers associated with memory (*Sell*) and NK-like (*Klrk1*, *Klrd1*) differentiation and downregulated genes related to exhaustion (*Lag3*, *Tigit*) and TCR signaling (*Cd3g*, *Cd3e*) (**Figure 6E**). In addition, exhaustion-prone clones upregulated gene signatures of cytokine production, T cell activation, inflammation, and cell adhesion and downregulated ribosome signature (**Figure 6F**). Thus, our results show that neoantigen-specific T cell clones with a lineage preference towards the T_SCM_ subset in the lymph node are less clonally expanded and less exhausted in the tumor.

**Figure 6.**
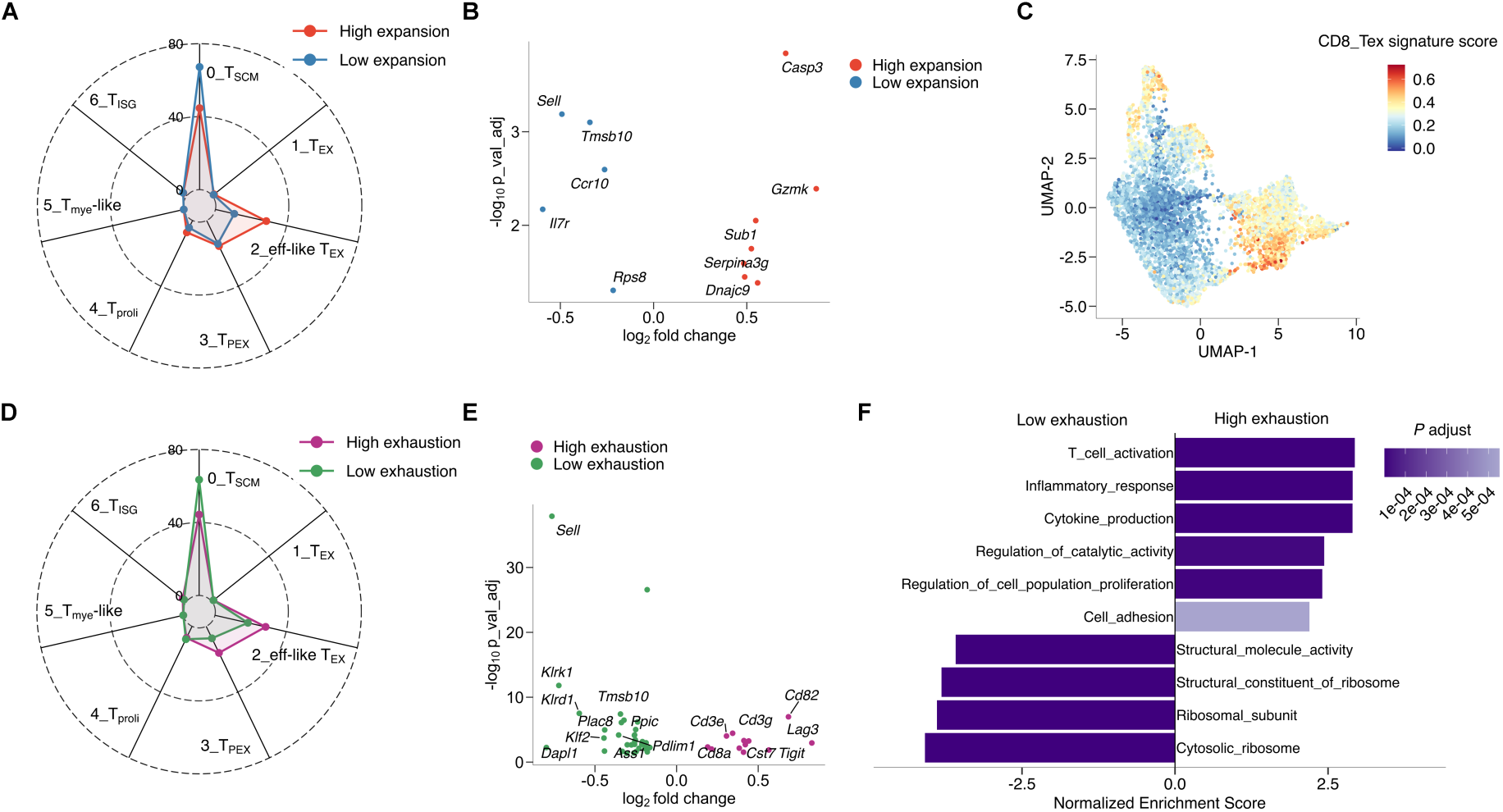
Differentiation traits of lymph node clones predict their transcriptional and clonal expansion states within tumor. (**A**) Radar plot showing the distribution of lymph node (LN) T cell subsets among SPAS-1 tetramer (Tet)^+^ CD8^+^ clones that exhibited high versus low clonal expansion in the tumor. (**B**) Volcano plot showing differentially expressed genes in LN Tet^+^ CD8^+^ clones with high versus low clonal expansion in the tumor. (**C**) UMAP plot illustrating the score of CD8_Tex signature at single-cell level. Cells are colored based on their CD8_Tex signature score. Data sourced from https://spica.unil.ch/refs/TIL/ ^61^. (**D**) Radar plot showing the distribution of LN T cell subsets among Tet^+^ CD8^+^ clones that exhibited high versus low exhaustion scores in the tumor. (**E**) Volcano plot showing differentially expressed genes in LN Tet^+^ CD8^+^ clones with high versus low exhaustion scores in the tumor. (**F**) GSEA comparing LN Tet^+^ CD8^+^ clones with high versus low exhaustion scores in the tumor. The x-axis represents the normalized enrichment score for each pathway, and the y-axis shows the corresponding pathways. The color-code indicates the adjusted *P* value.

### The cell fate decision between lymph node residence and tumor infiltration correlates with clinical outcomes in cancer patients

We sought to identify neoantigen-specific CD8^+^ T cell clones that showed different lineage preferences towards lymph node subsets versus tumor-infiltrating subsets. Lymph node (LN)- biased clones were more likely to differentiated into T_SCM_ cells, whereas TIL-biased clones were skewed towards T_PEX_ and eff-like T_EX_ subsets (**Figure 7A**). Compared to LN neoantigen specific CD8^+^ T cells from LN-biased clones, LN neoantigen specific CD8^+^ T cells from TIL-biased clones upregulated genes associated with exhaustion (*Tox*, *Tigit*, *Pdcd1*, *Lag3*) and effector function (*Gzmb*, *Gzmk*, *Nkg7*) and downregulated genes related to memory differentiation (*Il7r*) and interferon signaling (*Ifit3*) (**Figure 7B**). GSEA showed upregulation of chemokine, T cell activation and T cell apoptosis pathways in TIL-biased clones (**Figure 7C**). Next, we identified the gene signature of LN-biased neoantigen specific CD8^+^ TILs and that of TIL-biased neoantigen specific CD8^+^ TILs (**Figure 7D**). Notably, high expression of the TIL-biased gene signature in TILs correlated with disease-free survival in pan-cancer patients (**Figure 7E**). However, high expression of the TIL-biased gene signature negatively correlated with the response of melanoma patients to immune checkpoint inhibitors (**Figure 7F**). This result is consistent with the notion that lymph node serves as a reservoir for T cells to respond to immune checkpoint inhibitors^39,40,56^. Therefore, the lineage preference of T cell clones between lymph node and tumor-infiltration cell fates correlates with clinical outcomes in cancer patients.

**Figure 7.**
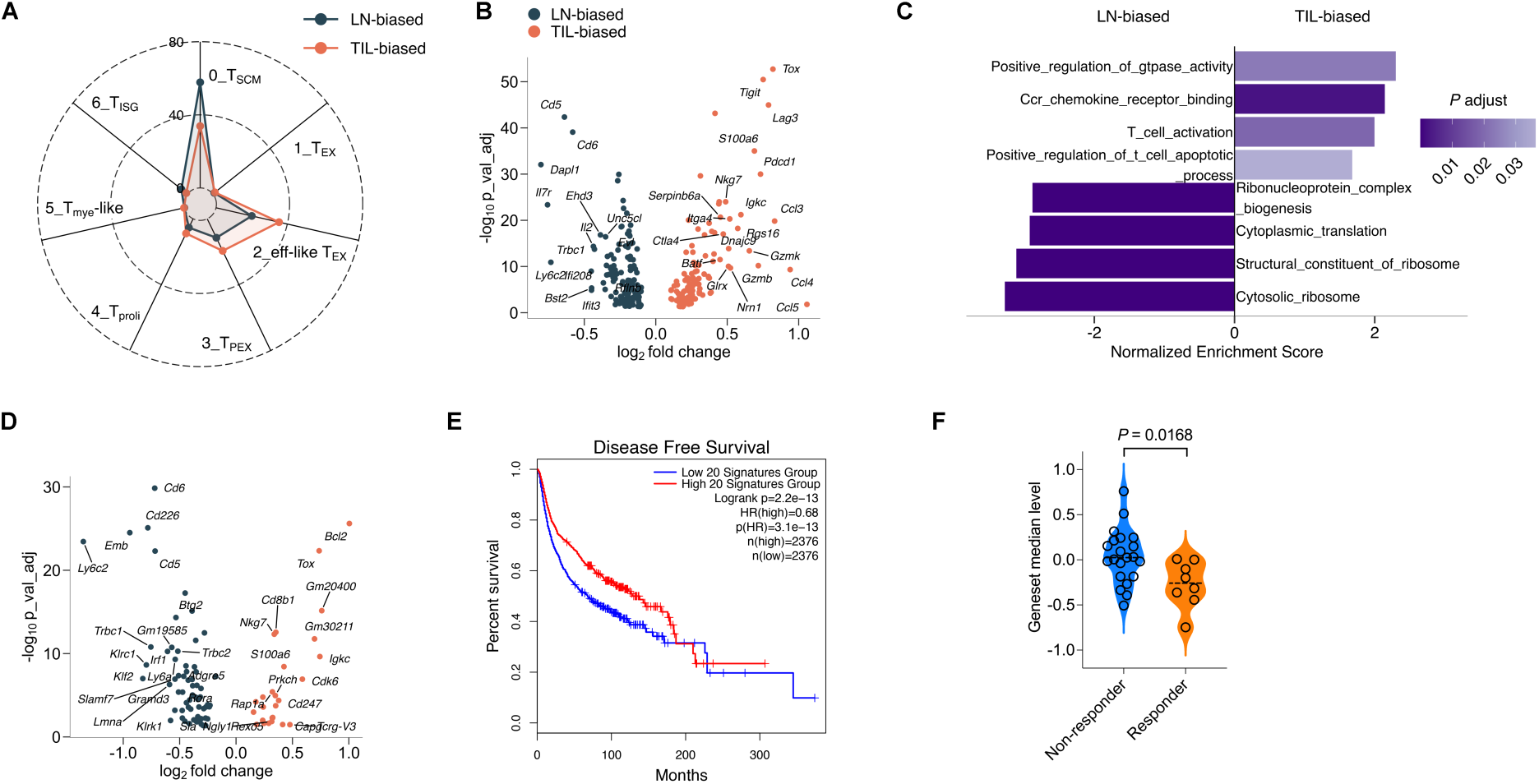
TIL-biased neoantigen-specific CD8^+^ clones exhibit a distinct transcriptional program that predicts clinical outcome in cancer patients. (**A**) Radar plot showing the distribution of lymph node (LN) T cell subsets among SPAS-1 tetramer (Tet)^+^ TIL-biased versus LN-biased CD8^+^ clones. (**B**) Volcano plot showing differentially expressed genes in LN Tet^+^ CD8^+^ T cells from TIL-biased versus LN-biased clones. (**C**) GSEA comparing LN Tet^+^ CD8^+^ T cells from TIL-biased versus LN-biased clones. The x-axis represents the enrichment score for each pathway, and the y-axis shows the corresponding pathways. The color-code indicates the adjusted *P* value. (**D**) Volcano plot showing differentially expressed genes in Tet^+^ CD8^+^ TILs from TIL-biased versus LN-biased clones. (**E**) Kaplan-Meier analysis of disease-free survival in pan-cancer patients grouped based on the expression level (high versus low) of the top 20 marker genes of Tet^+^ CD8^+^ TILs from TIL-biased clones, as indicated in panel D. Data source: http://gepia2.cancer-pku.cn/#survival (RRID: SCR_026154) ^62^ (**F**) Violin plot illustrating the expression level of the TIL-biased gene signature in tumor-infiltrating T cells from melanoma patients with or without clinical response to anti-PD1/CTLA4 treatment. Data source: https://resilience.ccr.cancer.gov/ ^63,64^.

## Discussion

In this study, we used a syngeneic prostate cancer model with a defined neoepitope generated during natural tumorigenesis and simultaneously profiled the single-cell transcriptome and TCR sequences of tumor-reactive CD8^+^ T cells. We defined the heterogeneous differentiation programs of neoantigen-specific CD8^+^ T cells in the tumor and draining lymph node. The shared TCR clones dominating the neoantigen-specific response in both the lymph node and tumor creates opportunity to perform clonal tracing analysis. We identified distinct signatures of divergent and lineage-biased clones and uncovered the differentiation trajectory from neoantigen-specific CD8^+^ subsets in the lymph node to those in the tumor. Notably, our results show that the differentiation program of neoantigen-specific CD8^+^ T cells in the draining lymph node implicates their clonal expansion and exhaustion in the tumor. In addition, lineage preference of neoantigen-specific CD8^+^ T cells between the lymph node-resident subsets versus the tumor-residence subsets correlates with clinical outcome in cancer patients.

Neoantigens are generated from mutations occur during tumorigenesis, which can be presented by MHC and recognized by T cells ^57^. As neoantigens are unique to tumors and absent in normal tissues, T cell clones specific for neoantigens bypass central tolerance and are critical for the antitumor efficacy of cancer immunotherapies ^57^. Therefore, determining the unique gene signature of neoantigen-specific CD8^+^ T cells is important for understanding their differentiation program, identifying neoantigen-specific CD8^+^ T cells and TCRs for adoptive cell therapy and uncovering biomarkers to predict the clinical outcome of cancer immunotherapies. scRNA-seq is instrumental for the effort to understand the transcriptional program and heterogeneity of antitumor T cells ^25,49–53^. However, neoantigen-specific T cells in patients are rare, diverse, and highly personalized^52^, which challenges the lineage tracing of the same clone in different tissues. Although single-cell omics were performed with murine T cells recognizing model antigens ectopically expressed by murine tumors^39,40^, it is unclear whether model antigen-specific T cells have the same differentiation program as T cells recognizing naturally derived neoantigens. The TRAMP-C2 model used in this study contains an immunodominant neoantigen, SPAS-1, derived from a natural point mutation that occurred during tumorigenesis^54^. Using an IFNγ reporter mouse strain and SPAS-1 tetramer, we generated scRNA+TCR-seq profiles of SPAS-1-specific CD8^+^ T cells in the tumor and draining lymph node, tetramer^-^ IFNγ- YFP^+^ PD1^+^ antitumor CD8^+^ TILs, and IFNγ-YFP^-^ PD1^-^ bystander TILs. Compared to other antitumor CD8^+^ TILs, neoantigen SPAS-1-specific CD8^+^ TILs upregulates genes associated with costimulation (*Tnfrsf9*, *Tnfrsf4*) as well as T cell activation, proliferation, and cytokine production pathways. This result indicates that neoantigen-specific CD8^+^ TILs may experience stronger TCR stimulation and have a hybrid phenotype of both activation and exhaustion. Therefore, combining 4-1BB and OX40 with known neoantigen-specific TIL markers such as PD1, TIGIT, and CD39^58^ may improve identification of neoantigen-specific TIL. Recent studies suggest that tumor draining lymph nodes not only contain the T_PEX_ subset but also a T_SCM_ population that is spared from the exhaustion program and responds more potently to ICI^38,41,56^. In addition to T_PEX_ and T_SCM_ subsets, we identified a highly clonally expanded eff-like T_EX_ subset in the tumor draining lymph node that may represent the transition state of neoantigen-specific CD8^+^ T cells ready to exit the lymph node and infiltrate the tumor. Given that lymph nodes are abundant with T cells expressing markers similar to T_PEX_ or T_SCM_ cells, the eff-like T_EX_ phenotype may be more likely to enrich with neoantigen-specific CD8^+^ clones in the tumor draining lymph node.

The shared dominant neoantigen-specific CD8^+^ clones between the tumor and draining lymph node provides an opportunity to track the differentiation and lineage choices of the same T cell clone. We found that the T_PEX_ subset showed a higher TCR diversity and lower clonal expansion than the T_SCM_, eff-like T_EX_, or T_EX_ subset. Pseudotime analysis suggest a T_PEX_ → T_SCM_ differentiation trajectory. Consistent with our observation, recent studies in the chronic viral infection show that antiviral T_PEX_ cells differentiate into memory cells in the absence of antigen ^59,60^. These findings raise the possibility that the T_PEX_, T_SCM_, and eff-like T_EX_ subsets are positioned in distinct regions of the tumor draining lymph node with different antigen abundances and types of antigen-presenting cells. Future studies with spatial omics may elucidate the niches of these neoantigen-specific CD8^+^ subsets. Our clonal lineage tracing showed that the divergent neoantigen-specific CD8^+^ clones are larger clones with greater clonal expansion whereas the clones biased towards the T_PEX_, T_SCM_, or T_EX_ subset were smaller with less clonal expansion. The clones with a greater tendency to differentiate into the T_SCM_ subset in the draining lymph node showed less clonal expansion and a lower level of exhaustion in the tumor. Consistently, the tumor-biased clones expressed higher levels of exhaustion related genes and lower levels of stem/memory associated genes. In addition, the gene signature of tumor-biased clones negatively correlated with the response to ICI, consistent with the notion that stem-like T cells are critical for the response to ICI treatment ^39,40,56^. It warrants future studies to determine whether the lineage bias and/or tissue bias of neoantigen-specific CD8^+^ T cells is determined by the TCR affinity.

## Materials and Methods

### Mice and tumor inoculation

Male IFNγ-YFP reporter (GREAT) mice (RRID: IMSR_JAX:017580) were purchased from the Jackson Laboratory. All mice were maintained in a specific-pathogen-free facility and were used for tumor inoculation at 6-8 weeks of age. Mice were housed with a 12-hour light/dark cycle. Room temperature was kept between 20 °C and 25 °C. Humidity was maintained between 30% and 70%. 10^6^ TRAMP-C2 tumor cells (RRID: CVCL_3615) were injected subcutaneously into the flank of each GREAT mouse. Mice were euthanized if the tumor diameter exceeded 2 cm. Throughout the study, no tumors reached this maximum allowable size. All procedures and experiments related to animals were performed according to the protocols approved by the Institutional Animal Care and Use Committee at the UT Southwestern Medical Center (UTSW, protocol 103162 and 103111).

### Tissue processing and single-cell suspension preparation

Single-cell suspensions from lymph nodes were prepared by manual dissociation. Tissue was minced and gently passed through a 100-µm-diameter strainer. Cells were washed, centrifuged, and resuspended in FACS buffer (phosphate-buffered saline (PBS) supplemented with 2% fetal bovine serum, 2 mM EDTA, and 1% penicillin/streptomycin). For TRAMP-C2 tumor dissociation, tissue was cut into small fragments and enzymatically digested in Liberase TL-containing buffer for 40 mins at 180 rpm. The digested tissues were then pushed through a 100-µm-diameter strainer. Mononuclear cells were isolated using density gradient centrifugation with 44% and 67% Percoll solutions. Following centrifugation, cells were washed with RPMI 1640 medium supplemented with 2% calf serum to remove residual Percoll. The isolated cells were then resuspended in FACS buffer for subsequent staining or sorting.

### Flow cytometry and cell sorting

The antibodies and dyes used for flow cytometry and cell sorting were as follows: anti-mouse CD8a PE (Biolegend, 53-6.7, 1:200, Cat# 100708, RRID: AB_312747), anti-mouse PD1 BV421 (Biolegend, RMP1-30, 1:200, Cat# 109121, RRID: AB_2687080), LIVE/DEAD™ Fixable Aqua Dead Cell Stain Kit (Thermo Fisher, 1:500, Cat# L34966), the class I tetramer, H-2Db SPAS1 (STHVNHLHC) APC (1:100, NIH Tetramer Core). Flow cytometry analysis and cell sorting were performed with a BD LSRFortessa (RRID: SCR_018655) and a BD FACSAria II (RRID: SCR_018934). Analysis was performed with FlowJo (v10.8.1, RRID: SCR_008520).

### Sample preparation for scRNA+TCR-seq

Single-cell RNA-seq and TCR libraries were generated using the Chromium Next GEM Single Cell 5’ Gene Expression and V(D)J Reagent Kits (10x Genomics) according to the manufacturer’s protocol. Briefly, freshly sorted cells were loaded onto the 10x Genomics Chromium Single Cell Controller to partition cells into GEMs. The generated gene expression and TCR libraries were sequenced on an Illumina HiSeq 3000 (RRID: SCR_016386).

### Data analysis for scRNA+TCR-seq

Fastq files of gene expression were aligned to mm10 reference genome using cellranger count function (Cell Ranger, v6.0.0, RRID: SCR_017344), which performs filtering, barcode counting and UMI counting. Downstream analysis was conducted using R (v4.3.2, RRID: SCR_001905) and Seurat (v4.4.0, RRID: SCR_016341). Samples were merged using the Merge function in Seurat. Cells with more than 5% mitochondrial gene content or in the top or bottom 2% for the number of detected genes were classified as outliers and excluded from further analysis. Cell cycle phase was assigned using the CellCycleScoring function. Data were normalized and scaled with NormalizeData and ScaleData functions, respectively. During scaling, the effects of cell cycle, total RNA counts, and mitochondrial gene percentage were regressed out using the parameter: vars.to.regress = c(’CC.Difference’, ‘nCount_RNA’, ‘percent.mt’). The top 2000 variable genes were identified using FindVariableFeatures function. TCR (^Tr.v) genes and Ig (^Ig.v) genes were excluded from the variable gene list for PCA analysis during clustering LN SPAS-1 tetramer^+^ and TIL SPAS-1 tetramer^+^ populations, as well as TIL SPAS-1 tetramer^+^ and TIL SPAS-1 tetramer^-^PD1^+^YFP^+^ populations, using RunPCA function. UMAP embeddings and neighbors were determined with the top 20 PCAs using RunUMAP and FindNeighbors functions, respectively. Clusters were identified using FindClusters function. A small fraction of low-quality cells (<1% of total cells) was removed post initial clustering. Marker genes were identified using FindAllMarkers or FindMarkers function for pairwise comparisons (parameters: min.pct = 0.1, logfc.threshold = 0.1).

TCR clonality analysis was performed using the scRepertoire R package (v2.2.1, RRID: SCR_025691). The filtered_contig_annotations output from Cell Ranger was imported into Seurat to create a list of TCR genes and CDR3 sequences by cell barcodes using the combineTCR function. Clonotypes were defined based on the amino acid sequence of the CDR3 regions of TCRα and TCRβ chains. Clone size levels were categorized as follows: Single (1 cell), Small (2-45 cells), Medium (46-522 cells), and Large (>= 523 cells).

### Trajectory analysis

Monocle 2 (v2.30.0, RRID: SCR_016339) was utilized to construct cellular trajectories and infer pseudotemporal ordering. Single-cell expression matrices were imported into Monocle 2, and top 10 marker genes in each cluster were selected for dimensionality reduction using DDRTree. Cells were ordered along pseudotime based on their transcriptional profiles, enabling the identification of key transitions during differentiation. Branch points in the trajectory were analyzed to uncover lineage-specific gene expression dynamics.

### Gene set enrichment analysis (GSEA)

GSEA was conducted using the clusterProfiler R package (v4.11.0, RRID: SCR_016884) to identify biologically relevant pathways and gene sets associated with the experimental conditions. Differential expression results were used to generate ranked gene lists based on log2 fold changes. Enrichment scores were computed using the Kolmogorov-Smirnov statistic, and significance was assessed through 1,000 permutations of the gene sets. Gene sets with adjust *P* < 0.05 and enrichmentScore > 0.3 were considered statistically significant. The analysis utilized the MSigDB database (v2024.1, RRID: SCR_016863) as the reference gene set collection, focusing on Gene Ontology (GO) pathways. Visualization of enriched pathways was performed using bar plots and enrichment maps to highlight key biological processes.

### Gene signature scoring

To quantify the activity of predefined gene signatures at the single-cell level, we employed the AddModuleScore function from the Seurat package (v4.4.0, RRID: SCR_016341). Gene sets for and “CD8_Tex” were derived from ProjecTILs ^61^. Specifically, marker genes for CD8_Tex cluster were identified using the FindAllMarkers function in Seurat (parameters: min.pct = 0.1, logfc.threshold = 0.1). Gene signature scores were calculated for each cell and visualized on UMAP embeddings to compare transcriptional activity across cell populations.

### Clone differentiation pattern analysis

For clone differentiation pattern analysis, clones with a size > 2 in tetramer^+^ cells were included. K-means clustering was performed on the percentage distribution of clones across seven predefined clusters derived from LN tetramer^+^ and TIL tetramer^+^ samples. To determine the optimal number of clusters (k), we calculated the total within-cluster sum of squares (WSS) for k ranging from 1 to 10. By evaluating the trend of WSS across different k values, we selected k = 4 as the number of clusters. This decision was informed by the distribution characteristics of the data and the analytical objectives, aiming to identify the major categories of T-cell clone differentiation patterns. The k-means algorithm was applied with 25 random starts to ensure robust clustering. The resulting clusters were used to classify clones into distinct differentiation patterns.

### Classification of clones for tissue-biased distribution, expansion, and exhaustion level analysis

To define TIL-biased and LN-biased clones, shared clones among TIL SPAS-1 tetramer^+^ and LN SPAS-1 tetramer^+^ populations were included. TIL-biased clones were identified as those with a modified percentage > 50% in TIL SPAS-1 tetramer^+^ population after normalization for population cell number. LN-biased clones were identified as those with a modified percentage > 50% in LN SPAS-1 tetramer^+^ population after normalizing population cell number.

To define the clones with high and low expansion, shared clones among TIL SPAS-1 tetramer^+^ and LN SPAS-1 tetramer^+^ populations were included. Clones with high expansion were classified as those with a clone size >= 6 in TIL SPAS-1 tetramer^+^ population, while clones with low expansion were classified as those with a clone size < 6 in TIL SPAS-1 tetramer^+^ population.

To define clones with high or low exhaustion levels, shared clones among TIL SPAS-1 tetramer^+^ and LN SPAS-1 tetramer^+^ populations were included. Clones with high exhaustion level were defined as those with a mean CD8_Tex signature score > 0.35 in TIL SPAS-1 tetramer^+^ population, while clones with low exhaustion level were defined as those with a mean CD8_Tex signature score <= 0.35.

### Statistics and reproducibility

Statistical analyses were conducted using R (v4.3.2, RRID: SCR_001905) and GraphPad Prism (v10.3.0, RRID: SCR_002798). Statistical significance was determined using a two-tailed unpaired t-test or a Chi-square test, with a significance threshold set at *P* < 0.05. Statistical significance in differential gene expression analysis was defined by an adjusted *P* < 0.05. All statistical parameters are detailed in the corresponding figures. Data distribution was assumed to be normal; however, formal normality testing was not performed. No data was excluded from the analysis.

## Data Availability

scRNA+ATAC-seq data in this study is publicly available in Gene Expression Omnibus (GEO) at GSE290142.

## Authors’ Contributions

**Y. Luo:** Data curation, formal analysis, investigation, methodology, writing. **C. Yao:** Data curation, formal analysis, investigation, methodology, writing. **T. Wu:** Conceptualization, data curation, formal analysis, investigation, methodology, supervision, funding acquisition, project administration, writing.

## Acknowledgments

We thank the following funding supports: AI158294 from National Institutes of Health, Clinic & Laboratory Integration Program from the Cancer Research Institute, and V Scholar Award from the V Foundation to T. Wu.

