## Supplemental Figures for "Single-cell clonal lineage tracing identifies the transcriptional program controlling the cell fate decisions by neoantigen-specific CD8^+^ T cells"

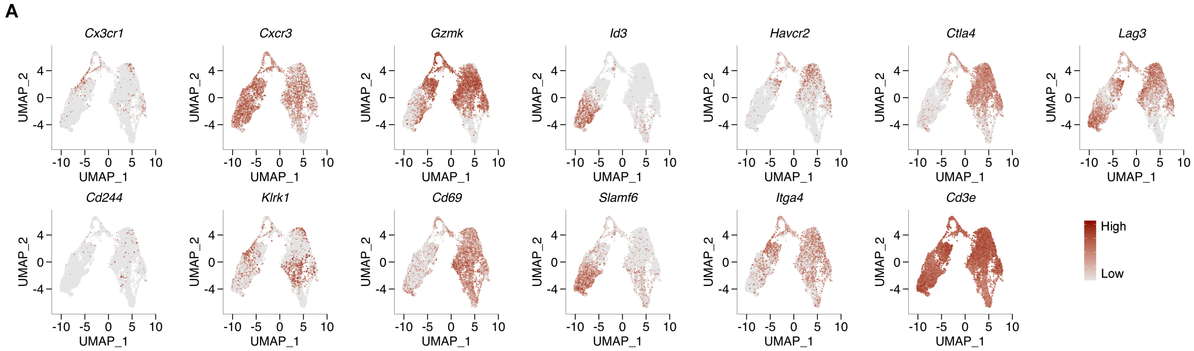


**Supplementary Figure 1**. Single-cell transcriptional profiling of selected genes among CD8$^+$ T subsets in the tumor-draining lymph node and the tumor. Gene expression levels are color-coded.

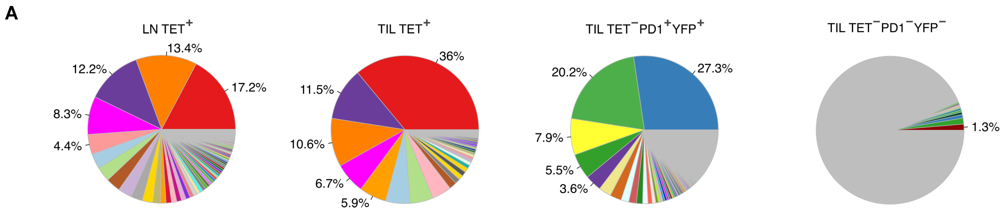


**Supplementary Figure 2**. Clonal analysis reveals the clonal dominance of neoantigen-specific $CD8^+$ T cells in the lymph node and tumor. (**A**) Pie charts illustrating the cell number proportion of unique clones across LN $Tet^+$, TIL $Tet^+$, TIL $Tet^-PD1^+IFN\gamma\text{-}YFP^+$, and TIL $Tet^-PD1^-IFN\gamma\text{-}YFP^-$ samples. Each color represents a unique clone.

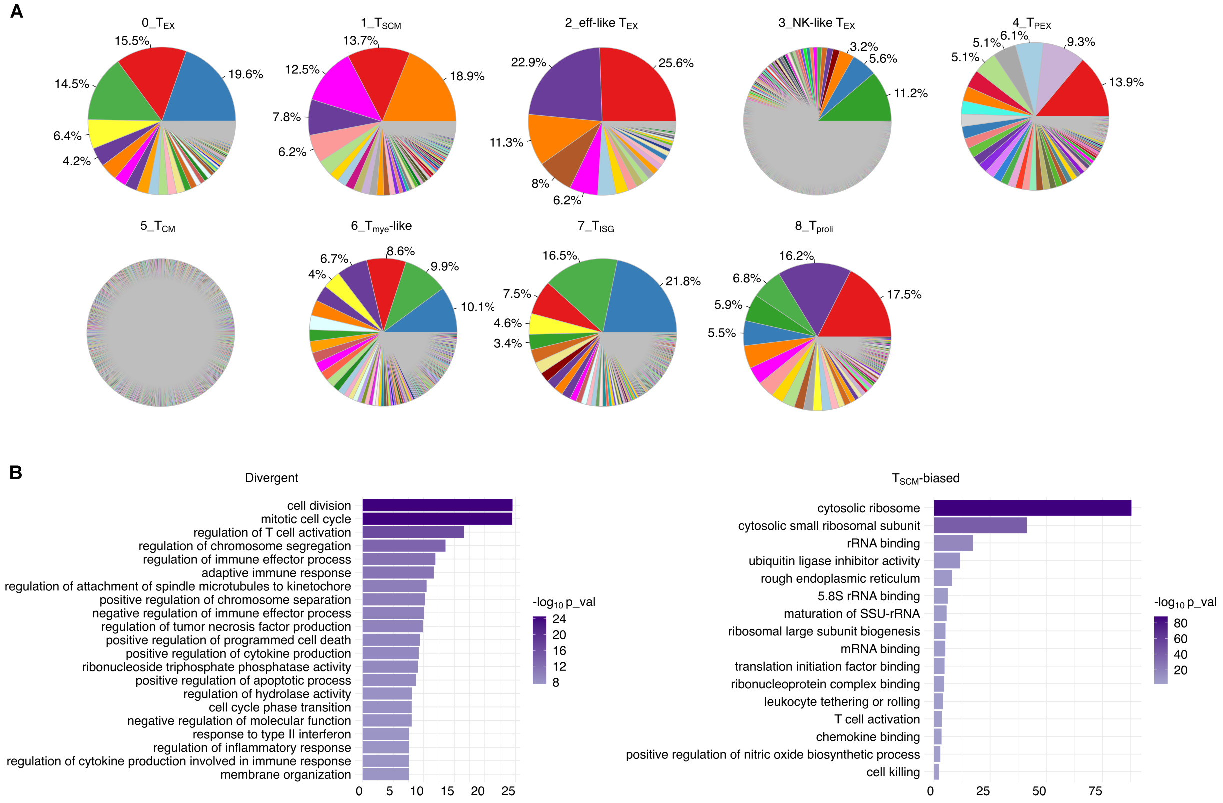


**Supplementary Figure 3**. Neoantigen-specific CD8$^{+}$ T cell clones in prostate cancer demonstrate divergent differentiation trajectories and distinct transcriptional biases. (**A**) Pie charts illustrating the clone proportions among different T cell subsets. Each color represents a unique clone. (**B**) Bar graphs depicting gene set enrichment analysis of marker genes from divergent and $T_{SCM}$ clones, generated with Metascape (RRID: SCR_016620). Data source: https://metascape.org/ [65]
